# Genome rearrangements drive evolution of ANK genes in *Wolbachia*

**DOI:** 10.1101/2023.10.25.563763

**Authors:** Ekaterina V. Vostokova, Natalia O. Dranenko, Mikhail S. Gelfand, Olga O. Bochkareva

## Abstract

**Introduction:** Genus *Wolbachia* comprises endosymbionts infecting many arthropods and nematodes; it is a model for studying symbiosis as its members feature numerous, diverse mutualistic and parasitic adaptations to different hosts. In contrast to nematode-infecting *Wolbachia,* genomes of arthropod-infecting strains contain a high fraction of repetitive elements creating possibilities for multiple recombination events and causing genome rearrangements. The mechanisms and role of these features are still not fully understood.

**Results:** Transposons cover up to 18% of an arthropod-infecting *Wolbachia* genome and drive numerous genome rearrangements including inversions and segmental amplifications. ANK (ankyrin-repeat domain family) genes are also often found at the breakpoints of rearrangements, while less than 7% of them were found within locally collinear blocks (LCBs). We observed a strong correlation between the number of ANK genes and the genome size as well as significant overrepresentation of transposons adjacent to these genes. We also revealed numerous cases of integration of transposases to the ANK genes affecting the sequences and putative products of the latter. Our results uncover the role of mobile elements in the amplification and diversification of ANK genes.

**Conclusions:** Evolution of arthropod-infecting *Wolbachia* was accompanied by diverse genome rearrangements driving the evolution of ANK genes important for bacteria-host interactions. This study demonstrates the effectiveness of our LCB-based approach to the *Wolbachia* genomics and provides a framework for understanding the impact of genome rearrangements on their rapid host adaptation.

## 1. Introduction

*Wolbachia* are obligate intracellular Gram-negative α-proteobactera of order Rickettsiales, family Anaplasmataceae. Genera *Ehrlichia*, *Anaplasma*, and *Rickettsia* are the closest to *Wolbachia*; however, the *Wolbachia* tree has not been rooted yet due to the absence of suitable, closely related organisms (Kaur et al. 2021). According to the conventional classification, *Wolbachia* strains are divided into at least 17 supergroups: A-F, H-Q, and S (Lo et al. 2002, Taylor et al. 2018, Laidoudi et al. 2020, Lefoulon et al. 2020). The majority of sequenced genomes belong to the A and B supergroups, while some supergroups are represented by only one genome.

*Wolbachia* are the most common intracellular symbiont, infecting a wide range of invertebrates from the superphylum *Ecdysozoa*, including filarial nematodes and arthropods such as insects, mites, and spiders (Zug and Hammerstein 2012, Weinert et al. 2015). According to published estimates, *Wolbachia* are prevalent in around 40-50% of insect species (Zug and Hammerstein 2012, Weinert et al. 2015). While mutualistic *Wolbachia* of nematodes are believed to be inherited strictly vertically, multiple host-switching events in arthropod *Wolbachia* suggest possible horizontal gene transfer and parallel adaptations to hosts (Lo et al. 2002).

*Wolbachia* infecting arthropods and nematodes have different effects on the hosts, ranging from reproductive manipulation to mutualism. Arthropod *Wolbachia* are mostly facultative to their hosts with some notable exceptions. In bedbugs, *Wolbachia* may provide essential B vitamins absent in the host diet (Hosokawa et al. 2010). Some arthropod-infecting *Wolbachia* strains possess a complete biotin operon, providing nutritional benefits to a host in a mutualistic manner (Balvín et al. 2018). Moreover, strains from bedbug *Cimex lectularius* and flea *Ctenocephalides felis* are essential for the host growth and reproduction, provisioning essential B vitamins (Nikoh et al. 2014, Driscoll et al. 2020). Similarly, if parasitic wasp *Asobara tabida* is treated with antibiotics, females cannot develop eggs (Dedeine et al. 2001). However, these are rare examples. On the other hand, many arthropod-infecting *Wolbachia* manipulate the host reproduction, increasing *Wolbachia* chances of transmission (Werren 1997). The *Wolbachia* infection can lead to cytoplasmic incompatibility, male killing, feminization, or parthenogenesis, and some genes involved in these processes are known; however, no strong connection between *Wolbachia* genome features and host phenotypes has been established so far (Werren 1997, 2021). Moreover, *Wolbachia* can affect the activity of the host’s transposable elements, depending on the host genotype, at least in *Drosophila melanogaster* (Eugénio et al. 2023). In contrast, in nematodes, *Wolbachia* are obligate essential symbionts, nutritional mutualists required for the development and long-term survival of onchocercid nematodes (Manoj et al. 2021). In *Brugia malayi* and *Onchocerca ochengi*, *Wolbachia* symbionts feature genes necessary for the heme metabolism and the synthesis of riboflavin (vitamin B2), both processes being essential for the host survival (Foster et al. 2005, Darby et al. 2012). *Wolbachia* may also contribute to the host nucleotide metabolism (Newton and Rice 2020).

The genome size of *Wolbachia* varies between 1.2 and 1.8 Mb in arthropod-parasitic species and between 0.96 and 1.1 Mb for mutualistic species of nematodes (Scholz et al. 2020). Clustering genes from 43 reference genomes of *Wolbachia* spp. at 80% threshold nucleotide identity resulted in 10,725 gene families (Scholz et al. 2020). Significant differences in the gene content were observed even within one supergroup. More functional acquisitions and depletions were found in supergroups C, D, and F, as opposed to supergroups A and B. The authors hypothesized that genomes in the C, D, and F supergroups belong to highly adapted *Wolbachia* strains with degraded genomes. Recent studies partly attribute the ability of *Wolbachia* to adapt to a wide range of arthropod hosts to segmental and single-gene duplications in genes encoding DNA methylases, bZIP transcription factors, and heat shock protein 90 (Liu et al. 2023).

A common feature of all insect *Wolbachia* genomes is a high fraction of repetitive elements such as insertion sequences, group II introns, duplicated segments of prophages, and multi-gene families (Chafee et al. 2010, Siozios et al. 2013, 2021). Insertion sequences cover approximately 10% of *Wolbachia* genomes and may play an important role in their evolution and adaptation (Cordaux et al. 2008, Kaur et al. 2017). Transposable elements belonging to the IS256 family disrupt the *wspB* gene (Sanogo, 2007) and might participate in the movement of *cifA* and *cifB* between or within *Wolbachia* genomes (Cooper et al. 2019). Group II introns, also frequent in *Wolbachia*, are supposed to be related to genome rearrangements and horizontal gene transfer (Leclercq et al. 2011). Reverse transcriptases originating in group II introns were found to stimulate chromosomal rearrangements, insertions, and duplications of prophages (Bordenstein and Bordenstein 2022). A diagnostic set of markers based on the IS site polymorphisms and genome rearrangements was developed for *Drosophila*-infecting *Wolbachia* (Kaur et al. 2017). The largest gene family present in *Wolbachia* strains infecting arthropods is the ankyrin-repeat domain (ANK) family highly prone to paralogization (Papafotiou et al. 2011, Siozios et al. 2013). ANK repeats are ∼33-residue sequence motifs devoid of enzymatic activity which mediate specific protein-protein interactions [(Mosavi et al. 2004)]. As ANK proteins might be secreted into the host cytoplasm to interact with host factors, they are supposed to play a major role in the *Wolbachia*-host interactions (Siozios et al. 2013).

Accumulation of genomic repeats causes substantial changes in the genome composition and gene order. Genome rearrangements occurring independently in different lineages may indicate parallel adaptation to new environments (Brandis and Hughes 2020, Dobrindt et al. 2010, Seferbekova et al. 2021). Moreover, reversible genome rearrangements in pathogens and symbionts may be responsible for the phenotype variety in populations via diverse mechanisms of phase variation, which are common for multi-gene families involved in bacteria-host interaction (Kojic et al. 2020, Cui et al. 2012, Shelyakin et al. 2019). Recently we developed a LCB-based approach to study genome rearrangements in closely related bacterial strains (Seferbekova et al. 2021, Zabelkin et al. 2022). Here we demonstrate the effectiveness of this approach to *Wolbachia* genomics.

Here, we provide a comprehensive analysis of available complete genomes of *Wolbachia* focusing on genomic changes responsible for the host adaptation. Firstly, we analyze the pangenome composition and host-specific gene families. Then, we study the composition and expansion rates of repetitive elements focusing on the interplay between IS families and ANK genes. Finally, we reconstruct the history of genome rearrangements revealing parallel events such as inversions and segmental amplifications.

## 2. Materials and Methods

### Dataset

We collected 159 complete *Wolbachia* genomes available in the RefSeq database as of March 2023 **(Supplementary Table 1)**. The host information was obtained from the assembly source metadata.

### Genome analysis using the PanACoTA pipeline

`PanACoTA` is a toolbox for large-scale comparative genomics of bacteria (Perrin and Rocha 2021). The pipeline contains six modules, which were run successively with the following parameters: at step *`prepare`* only complete genomes were included; step *`annotate`* was performed using `Prokka` v1.14.6 (Seemann 2014) and `Prodigal` v2.6.3 (Hyatt et al. 2010); step *`pangenome`* was performed with 60% protein similarity threshold; step *`corepers`* was used to select single-copy universal genes; step *`align`* was performed using `MAFFT` v7.520 (Katoh and Standley 2013); step *`tree`* inferred a phylogenetic tree from the concatenation of alignments of single-copy universal genes using `IQTree` version 2.2.0.3 (Minh et al. 2020) with 1000 bootstraps and the GTR DNA substitution model. The genomes of *Rickettsia conorii* str. Malish 7, *Rickettsia prowazekii* str. Breinl, *Ehrlichia ruminantium* str. Welgevonden were used as outgroups. The tree was visualized with `iTOL` version 6.7.4 (Letunic and Bork 2021).

### Construction of the pan-genome

Orthology groups were constructed using `ProteinOrtho` V5.13 (Lechner et al. 2011) with parameters *cov*=50 (at least 50% coverage of both proteins in `BLAST` alignments) and *identity*=60 (at least 60% identity in common segments).

### Reconstruction of gene losses and gains

Gene gains and losses was reconstructed using `CAFE 5` version 1.1 (Mendes et al. 2021) which takes in a phylogenetic tree and the counts of orthologous groups as input.

### Annotation of ISs

We used two strategies to annotate insertion sequences; `ISEScan` v. 1.7.2.3 (Xie and Tang 2017) and `ISFinder` (Siguier et al. 2006) with subsequent manual check of genome coordinates in NCBI. `ISEScan` predicted more regions with insertion sequences, but some of them overlapped single-copy house-keeping genes, and hence the results of `ISFinder` were used for the downstream analyses.

### Reconstruction of locally collinear blocks

For the initial assessment of rearrangements, dot plots were built using `D-GENIES` version 1.4 (Cabanettes and Klopp 2018) for the pairwise comparison of genomes. Locally collinear blocks (LCBs) were constructed with `SibeliaZ` version 1.2.5 (Minkin and Medvedev 2020). Optimal parameters for the blocks reconstruction were calculated using the badlon 0.1.3 *prepare* module (-*k* 15 -*a* 1740 -*n* for clade 1; -*k* 15 -*a* 1160 for clade 2) (Zabelkin & Bochkareva, in preparation). The minimal length of blocks was set to 500bp. LCB quality was estimated using the `badlon` 0.1.3 *`analysis`* module. The blocks were annotated using the `badlon` 0.1.3 *`annotate`* module.

### Estimation of rearrangement frequency

For this LCBs were constructed for each genome pair separately. Mash distances generated during the PanACoTA *prepare* step were used, and genome pairs were filtered so that each genome pair was included only once. Given the number of LCBs for each genome pair and the coverage, the rearrangement frequency was estimated and visualized using the `seaborn Python` library.

### Reconstruction and analysis of genome rearrangements

Genome rearrangements were analyzed using `PaReBrick` v.0.5.5 which takes LCBs to reconstruct the rearrangement events according to a given phylogenetic tree (Zabelkin et al. 2022). Two types of genome rearrangements were considered, balanced ones which change only the block order (inversions and translocations) and unbalanced ones which change the block copy number (insertions, deletions, and duplications).

### Prediction and alignment of ANK proteins

To find genes encoding members of ankyrin repeat (ANK) multi-family, we first searched for proteins with ankyrin repeat domains by `PfamScan.pl` (Finn et al. 2014) with PF00023.33, PF12796.10, PF13606.9, PF13637.9, and PF13857.9 `PFAM` entries and then performed a `BLASTP` search (Camacho et al. 2009) for all ORFs including the short ones with 90% identity and coverage thresholds. Multiple alignment of ANK multi-family genes was done using `muscle` (Edgar 2004).

## 3. Results

### Genome size, pangenome, and phylogeny

We studied 159 complete *Wolbachia* genomes from RefSeq **(Figure 1A)**. Two thirds of genomes were from hosts belong to three insect orders (Diptera – 64 genomes; Lepidoptera – 47 genomes; Hymenoptera – 19 genomes); ten *Wolbachia* genomes from nematodes; one, from Collembola; one, from Araneae.

**Figure 1.**
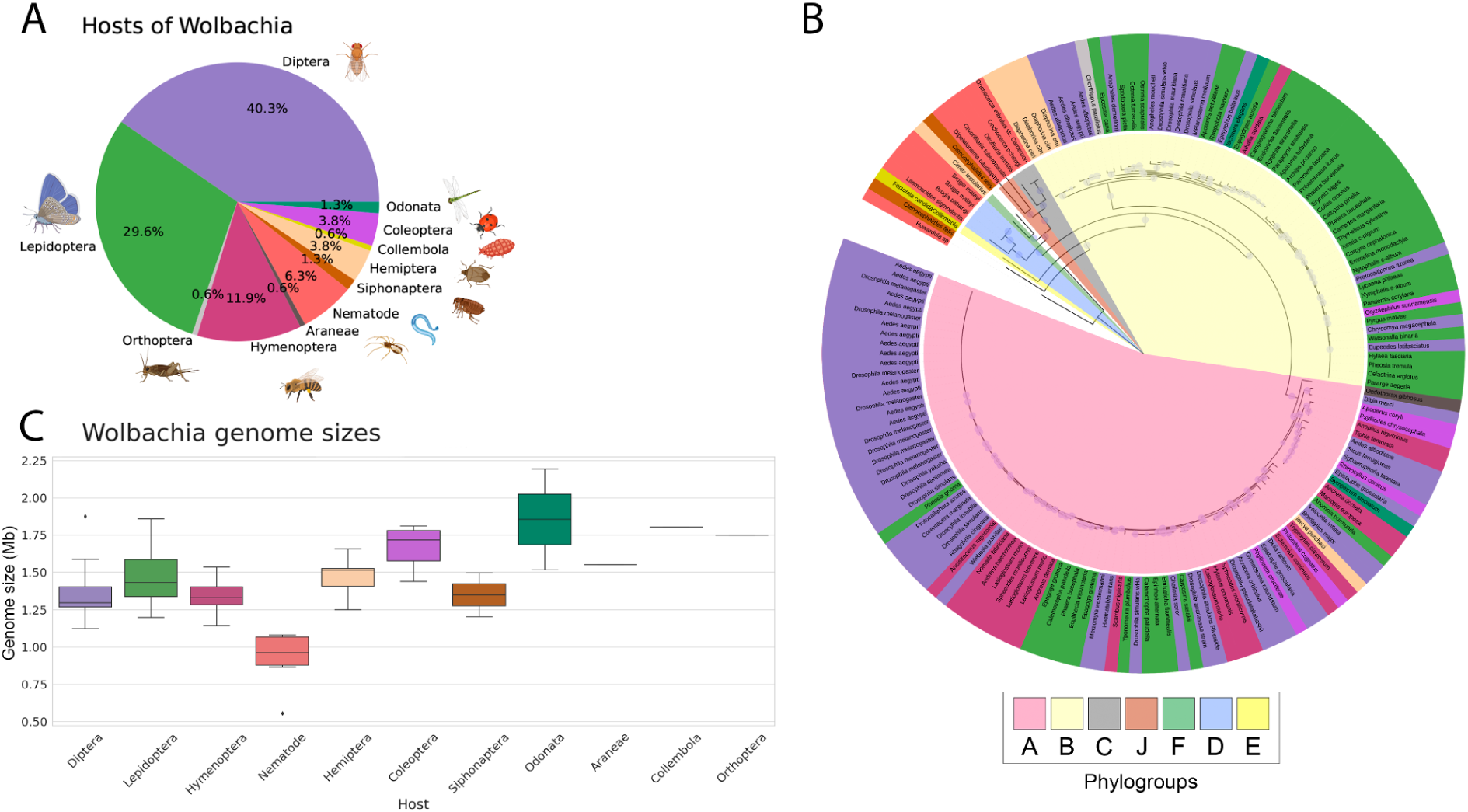
*Wolbachia* genomes and their hosts. **(A)** Composition of the RefSeq *Wolbachia* dataset of completely 159 assembled *Wolbachia* genomes; **(B)** Rooted phylogenetic tree of *Wolbachia* genomes. **(C)** Genome size of *Wolbachia* from different hosts. Сolors represent the host taxonomy in (A), (B), and the outer ring of (C); colors in the internal part of (C) indicate the supergroup, see the legend.

The pangenome of 159 *Wolbachia* strains was constructed with the 60% amino acid similarity threshold. It contained 14893 orthologous groups and 9193 singletons. The U-curve featured 312 *core* (universal) genes, a large *cloud* fraction comprising genes present in few genomes, and a limited *shell* fraction of genes found in five or more, but not all strains **(Supplementary Figure 1)**. The U-curve is homogenous without distinct internal peaks, which indicates the absence of large cohorts of genes specific to a distinct clade or group of genomes. In agreement with the U-curve, the fractional composition graph demonstrates that the *Wolbachia* pangenome is open showing no saturation due to a large fraction of singletons; the core genome also does not reach the plateau **(Supplementary Figure 2)**. All other fractions stabilize quickly.

The maximum likelihood phylogenetic tree was constructed based on concatenated alignment of 192 single-copy core genes with 1000 bootstraps **(Figure 1B)**. As expected, *Wolbachia* from arthropods form two large clades which are supergroups A and B in the conventional classification (Scholz et al. 2020). Nematode-infecting *Wolbachia* form a monophyletic clade that also includes several *Wolbachia* from arthropod hosts, such as *Cimex lectularius*, *Folsomia candida,* and *Ctenocephalides felis* (cat flea). The tree topology does not agree with that of the hosts indicating multiple, parallel reinfections of arthropod species.

Note that nematode-infecting *Wolbachia* have significantly smaller genomes (0.9-1.1 Mb) than others, including arthropod-infecting strains from the same branch (1.2-1.8 Mb, **Supplementary Figure 3**). The largest genome (2.2 Mb) belongs to *Wolbachia* sp. from Odonata, *Ischnura elegans* **(Figure 1C)**.

### Gene gains and losses

We estimated the rates of gene gains and losses according to the constructed phylogeny **(Figure 2A,B)**. Both rates were higher in leaves compared to internal nodes, but even for them, the number of events was quite low (maximum 10-20 genes lost or gained). Thus, we did not observe large clade-specific gene gains or losses, in agreement with the pan-genome U-curve analysis.

**Figure 2.**
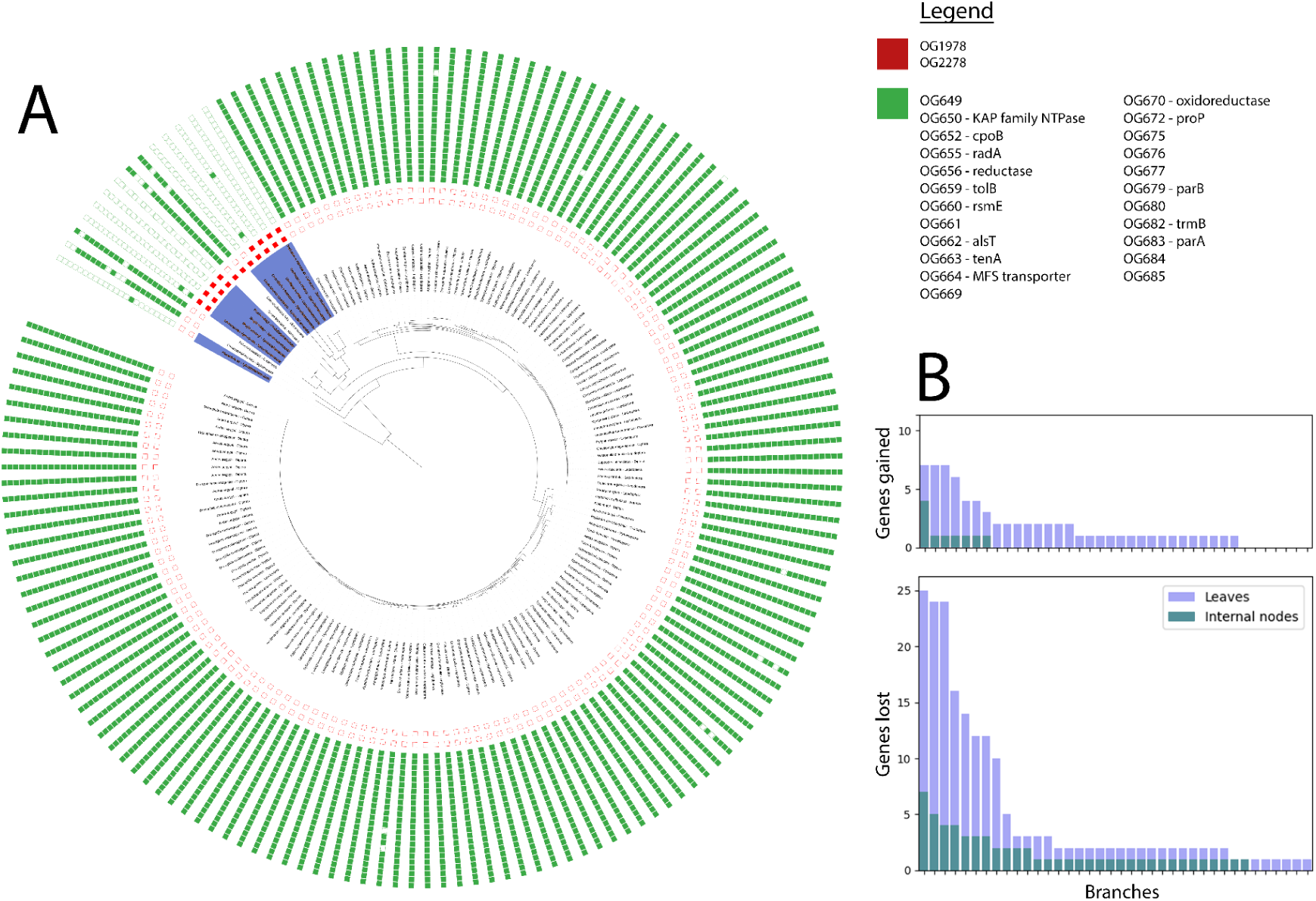
Gene gains and losses in *Wolbachia*. The number of genes **(A)** lost and **(B)** gained for various species (leaves) or ancestral populations (internal nodes). **(С)** Phyletic patterns of the genes which have been gained (red) and lost (green) in nematode-infecting *Wolbachia* (highlighted by blue). The numbers of orthogroups are listed in the legend in the outward direction.

To analyze host-specific adaptation in *Wolbachia*, we searched for gene families which were present only in strains from particular hosts. In nematode-infecting *Wolbachia*, we observed the absence of a large set of genes including genes encoding DNA repair protein RadA, Tol-Pal system protein TolB, cell division coordinator CpoB, ribosomal RNA small subunit methyltransferase RsmE, a transcriptional regulator from the TenA family, MFS transporter **(Figure 2C)**. In turn, two genes encoding hypothetical proteins are present in nematode-infecting *Wolbachia* and only two arthropod-infecting strains, however, the phyletic patterns of these genes are consistent with single acquisition events and thus are less likely to be linked to host adaptation mechanisms. For other hosts, we did not observe host-specific gene gains or losses.

### Transposon and ANK gene composition

Overall, thirteen IS families were found **(Figure 3A)**. Phyletic patterns for some of them (IS3, IS6) are consistent with the tree topology, while for others (such as IS5) they are not. Interestingly, IS6 was found only in phylogroup B. We observed transposon bursts in various genomes, but they are not consistent with the tree structure or with host shifts.

**Figure 3.**
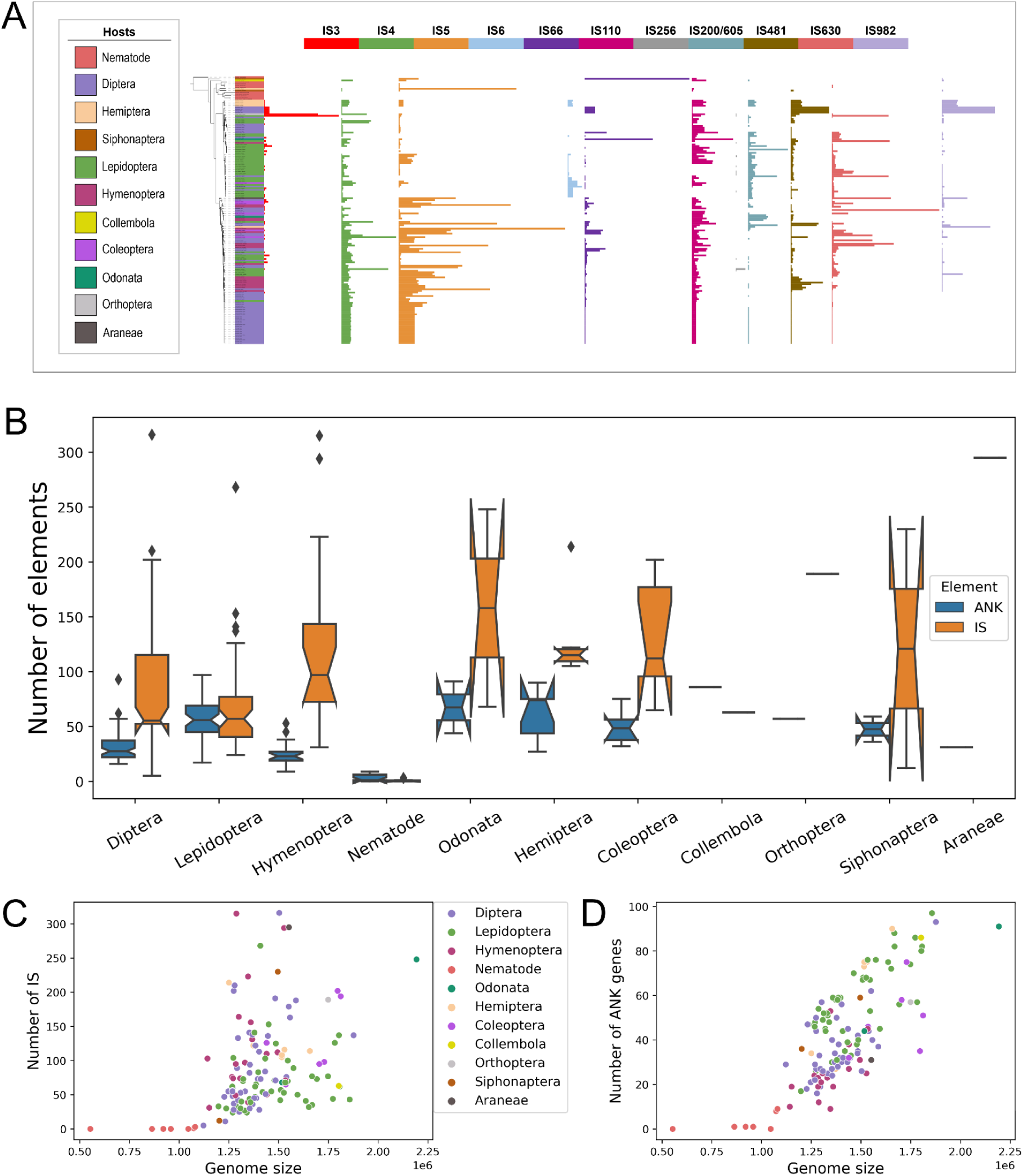
ISs in Wolbachia genomes. **(A)** IS copy number across the phylogenetic tree, colors reflect the IS families. **(B)** Number of ISs and ANK genes in *Wolbachia* genomes from different hosts. **(C)** Relationships between the number of ISs in a genome and the genome size. **(D)** Correlation between the number of ANK genes in a genome and the genome size.

In total, transposons cover up to 18% of a *Wolbachia* genome **(Supplementary Figure 4).** A low fraction of transposons was observed in nematode-infecting *Wolbachia*, which have significantly smaller genomes compared to arthropod-infecting strains; the highest, in *Wolbachia* with the largest genome from *Ischnura elegans* **(Figure 3B,C)**. Surprisingly, ANK genes were also almost absent in nematode-infecting *Wolbachia*; in turn, in arthropod-infecting strains we found up to 97 ORFs with at least one ANK-repeat domain (and up to 123 ORFs with BLAST hits to known ANK genes). Interestingly, we observed a strong correlation between the number of ANK genes and the genome size (Pearson’s *r* = 0. 78, *p* = 3.2 × 10^−33^) (**Figure 3B,D**).

Note that arthropod-infecting *Wolbachia* belonging to the nematode-infecting clade also have a relatively high fraction of ISs and large genome size. This allowed us to hypothesize that mobile elements may act as a driver of the ANK gene amplification. To check this, we estimated co-localization of ANK genes and transposons. Indeed, transposons were observed in a neighborhood of ANK genes significantly more often than expected, the null expectation calculated based on portions of transposons in the genomes (95% confidence intervals for the means were 0.15-0.17 and 0.05-0.06, the Wilcoxon test *p* = 1. 21 × 10^−26^)

### Rates of genome rearrangements

To estimate the rates of genome rearrangements we build the following model. The number of locally collinear blocks (and thus breakpoints between them) was considered as a proxy for the rearrangement frequency, and the phylogenetic distance between genomes was considered as the mutation frequency (see Methods). Then these values were estimated for all pairs of *Wolbachia* genomes. Note that the number of LCBs depends on the quality of alignment and hence decreases with the increasing distance between strains, thus this model has limitations.

Indeed, the number of LCBs and the distance between the genomes were positively correlated at evolutionary distances <0.06 substitutions per site (Pearson’s *r* = 0.65, *p* = 6.0 × 10^−249^) **(Supplementary Figure 5)**. In this range the genome coverage by constructed LCBs was high (>80%), thus the number of blocks indeed reflected the number of genome rearrangements between the genomes. At larger evolutionary distances, the genome coverage by blocks dropped dramatically and the representation of genomes by locally collinear blocks was not reliable. Hence, we analyzed genome rearrangements in the main *Wolbachia* clades separately.

### Parallel rearrangements and recombination hotspots

Rearrangements that occur several times independently could indicate selection pressure and thus be responsible for host adaptation. We focused on such parallel events and studied their occurrence in genomes from different hosts. For phylogroup A, 1016 locally collinear blocks of minimal length 500 bp were obtained providing 55% mean genome coverage. Reconstruction of rearrangements across the clade revealed 61 rearrangement hotspots which were involved in several independent inversions. In phylogroup B, 880 locally collinear blocks provide 59% mean genome coverage, the reconstruction yielded 21 rearrangement hotspots.

In both clades we observed no inversions correlated with the host phyletic pattern, although the number of parallel events was rather high. For example, the hotspot with the highest parallelism score (see Methods in (Zabelkin et al. 2022)) in phylogroup A is a ∼1200 kb region involved in five different inversions, the most common one of ∼50 kb length, and occurring at least six times independently in strains from different hosts **(Fig. 4A)**. The boundaries of the inversion were formed by IS3 family transposases on one side adjacent to an ANK gene. To systematically find genomic features responsible for genome rearrangements in *Wolbachia,* we searched for repetitive elements in the inversion breakpoints. In most cases transposons were found, which was expected considering the number of transposons in *Wolbachia* genomes. We hypothesized that the impact of different types of transposons on the rates of rearrangements could depend on their density in the genomes. However, we did not not observe any correlation between the number of cases when an IS element was observed at the inversion breakpoints and its average frequency of occurrence in genomes (data not shown).

**Figure 4.**
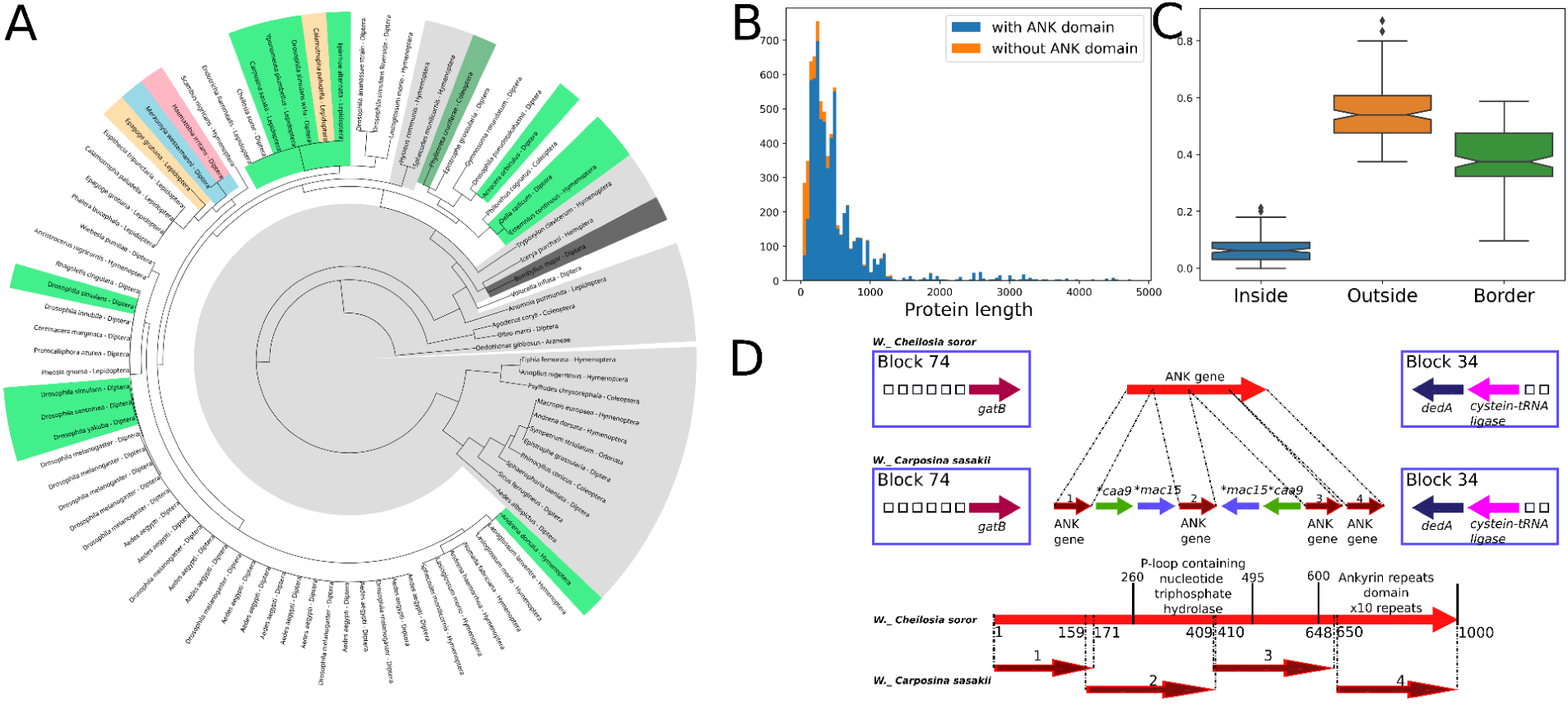
Genome rearrangements in Wolbachia. **(A)** An example of parallel inversions in phylogroup A. The tree is colored to reflect the states of the rearrangement hotspot comprising IS3 family transposases and an ANK gene. White represents the most common state (assuming no rearrangements), other colors reflect various inversions, gray represents complex rearrangements or series of inversions. A parallel inversion with the highest parallelism score (light green) occurred between two genomic repeats of IS3 transposases. **(B)** Fractions of ANK-encoding ORFs observed within LCBs and partly or fully outside LCBs. All pairwise comparisons are statistically significant. **(C)** Lengths of products of ORFs with (blue) and without (orange) ANK domain producing significant blast hits to ANK-encoding proteins. **(D)** An example of disruption of an ANK gene by an inverted repeat of IS5 family transposases Caa9 and Mac15 resulting in formation of four short ORFs in *Wolbachia* endosymbiont of *Carposina sasakii*. Arrow colors indicate genes from different orthogroups.

Consistent to our previous observations, ANK genes were also often found at the breakpoints of rearrangements. On average only 6.5% of ANK genes in a genome were found inside locally collinear blocks **(Fig. 4B)** which might be explained by their adjacency to transposons. In turn, transposon expansion and subsequent genome rearrangements may drive the fast evolution of ANK genes. To validate this hypothesis, we studied the composition of ANK genes in the genomes using blast search to proteins with the ANK domain (see Methods). We found 6286 ORFs containing the ANK domain and 759 additional ORFs with significant similarity to them in other domains. In total, 7.045 proteins were clustered in 624 orthogroups with the identity threshold of 60% **(Supplementary Figure 6A)**. Surprisingly, almost half of them were singletons, with others belonging to the accessory fraction of pan-genome; the largest orthogroup contains genes from only 96% of arthropod-infecting *Wolbachia* **(Supplementary Figure 6B)**. The length of gene products varied from 32 to 4793 amino acids **(Fig. 4C)**. Altogether, these observations indicate an exceptional variety of ANK genes.

To reveal the impact of insertion sequences on the domain structure of the encoded proteins, we aligned ANK genes from homologous loci defined as regions between two conserved CLBs in closely related *Wolbachia*. Indeed, we observed numerous cases of integration of transposases in ANK genes, affecting the sequences and putative products of the latter. An example of such integration is disruption of an ∼1 kb ANK gene, located in the genomic locus between genes encoding glutamyltRNA(Gln) amidotransferase GatB and cysteine-tRNA ligase, by IS5 family transposases observed in *Wolbachia* endosymbiont of *Carposina sasakii.* This resulted in formation of four short ANK ORFs of the lengths 171 nt, 252 nt, 240 nt, and 351 nt, all putatively transcribed **(Fig. 4D)**. The second ORF is surrounded by inverted repeats of transposases which may putatively act as a phase variation mechanism affecting the domain composition of the product.

### Segmental duplications

As expected, numerous segmental duplications were observed, affecting genes related to various metabolic pathways. In both phylogroups, the genome region with the highest copy number variation comprises genes which encode bifunctional protein GlmU, NAD(P)-dependent oxidoreductase, glycosyltransferase, a phytanoyl-CoA dioxygenase family protein, ABC transporter ATP-binding protein, GntG-family PLP-dependent aldolase, protein from the threonine aldolase family, MFS transporter, a protein from the UDP-glucose dehydrogenase family, and a DMT-family transporter. This ∼15 kb fragment has up to five copies per genome located in three distant loci (other copies are in tandem) which are also affected by smaller rearrangements such as gene duplications or deletions **(Supplementary Figure 7A)**. The gene trees demonstrate both tandem duplications in strains and horizontal gene transfer of the gene cassette in arthropods-infecting *Wolbachia* **(Supplementary Figure 7B)**. Moreover, this cassette is absent in nematode-infecting *Wolbachia* which is consistent with the hypothesis that these genes play a role in the adaptation to arthropods.

## 4. Discussion

Although *Wolbachia* have been discovered almost a century ago, they remain an object of intensive research as the genomic factors of host adaptation are still not clear. The number of complete *Wolbachia* genomes doubled in 2023, now comprising 159 genomes from different hosts. The structure of their phylogenetic tree **(Fig. 1C)** with multiple, parallel reinfections of arthropod species in A and B supergroups provides an opportunity to study events in different *Wolbachia* lineages using our novel computation approach (Seferbekova et al. 2021, Zabelkin et al. 2022).

Consistent to previous observations, nematode-infecting *Wolbachia* have smaller genomes with a low fraction of repetitive elements, such as transposons and ANK genes, compared to arthropod-infecting strains even from the same branch. Many other genes including *radA*, *tolB*, and *rsmE* are absent in nematode-infecting *Wolbachia* in comparison to arthropod-infecting strains. While many significant differences in the gene content were reported on smaller datasets (Scholz et al. 2020), we observed no host-specific gene gains or losses within arthropod-infecting clades. In turn, numerous segmental duplications affecting genes related to various metabolic pathways were found, which is consistent with a recent study suggesting that segmental and single-gene duplications play a role in the ability of *Wolbachia* to adapt to a wide range of arthropod hosts (Liu et al. 2023).

The rates of inversions in arthropod-infecting *Wolbachia* is also extremely high which agrees with the high number of repetitive events in the genomes. Indeed, transition to the intracellular lifestyle is often accompanied by accumulation of mobile elements causing substantial changes in the genome composition and gene order due to a more relaxed selective pressure (Mamirova et al. 2007, Seferbekova et al. 2021). The frequency of rearrangements in fast-evolving pathogens with a high number of mobile elements correlates with the presence of repeats (Darling et al. 2008, Bochkareva et al. 2018). Indeed, in the boundaries of inversions we observed transposons, covering up to 18% of an arthropod-infecting strains, which indicates their major role in genome instability of *Wolbachia*. We hypothesized that the impact of different types of transposons on the rates of rearrangements could depend on their density in the genomes, but observed no such correlation. This may be explained by fast evolution and pseudogenization of mobile elements or purifying selection acting on the rearrangements.

ANK genes were also often found in the breakpoints of rearrangements. Moreover, we observed a strong correlation between the number of ANK genes and the genome size as well as significant overrepresentation of transposons adjacent to them. This indicates that mobile elements may drive the ANK gene amplification and diversification. Previously, a comparison of representative strains from *Wolbachia* phylogroups A and B revealed that ANK genes demonstrated high sequence variability and were affected by intragenic recombination and horizontal gene transfer between strains (Siozios et al. 2013). This results in major changes in the encoded proteins, such as motif deletions, amino acid insertions, and also pseudogenization due to insertion of transposable elements conferring premature stop codons (Iturbe-Ormaetxe et al. 2005, Siozios et al. 2013). Using a novel LCB-based approach, we demonstrated genetic variability of ANK genes even in closely related strains, due to integration of ISs and genome rearrangements. They not only significantly expand the ANK repertoire in *Wolbachia* but may affect the level of gene expression and domain composition of their products via inversions between inverted repeats.

Most of the observed structural variants were not monophyletic, but were present in phylogenetically distant genomes. Although we did not observe any host-specific rearrangements, the number of parallel events was very high, including large-scale reversible inversions and segmental duplications of gene cassettes involved in the metabolism. Altogether, the observed parallel rearrangements may indicate phenotypic variation allowing *Wolbachia* strains to adapt rapidly to new environmental conditions.

Accumulation of genomic repeats in arthropod-infecting *Wolbachia* drives multiple recombination events which may contribute to host specialization and enable rapid adaptation. This study demonstrated the effectiveness of the LCB-based approach to *Wolbachia* genomic data and expanded our understanding of the *Wolbachia* evolution. Long-read sequencing of *Wolbachia* accompanied by transcriptomic analysis would shed light on connections between large-scale variations in *Wolbachia* genomes, such as inversions, gene and segmental duplications, composition of multi-gene families, and the host phenotypes.

## Author Contributions

Conceptualization, MSG and OOB; methodology, OOB; analysis, EVV and NOD; writing - original draft preparation, EVV and OOB; writing - review & editing, MSG; visualization, EVV and NOD; funding acquisition, MSG and OOB. All authors have read and agreed to the published version of the manuscript.

## Data Availability Statement

The datasets supporting the conclusions of this article and used ad hoc scripts are available via the link https://github.com/OlgaBochkaryova/wolbachia-genomics

## Acknowledgements

We thank Ekaterina Kolodyazhnaya and Alexey Zabelkin for assistance with the Parebrick and Badlon tools.

## Supplementary Materials

**Table S1**: List of complete *Wolbachia* genomes with host information and composition of insertion sequences.

**Figure S1:**
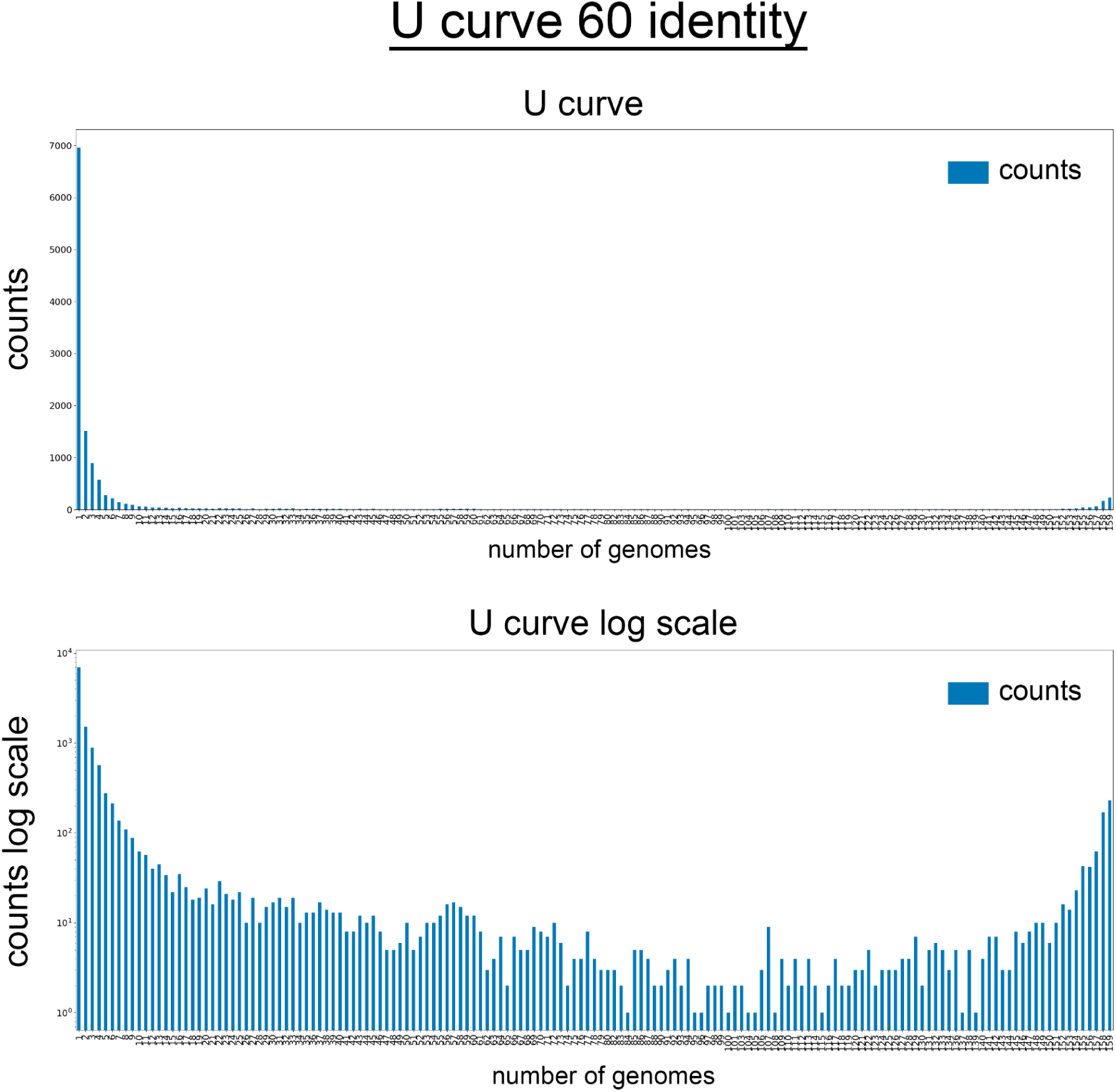
Distribution of orthogroups by the number of strains in which they are present (U-curve).

**Figure S2:**
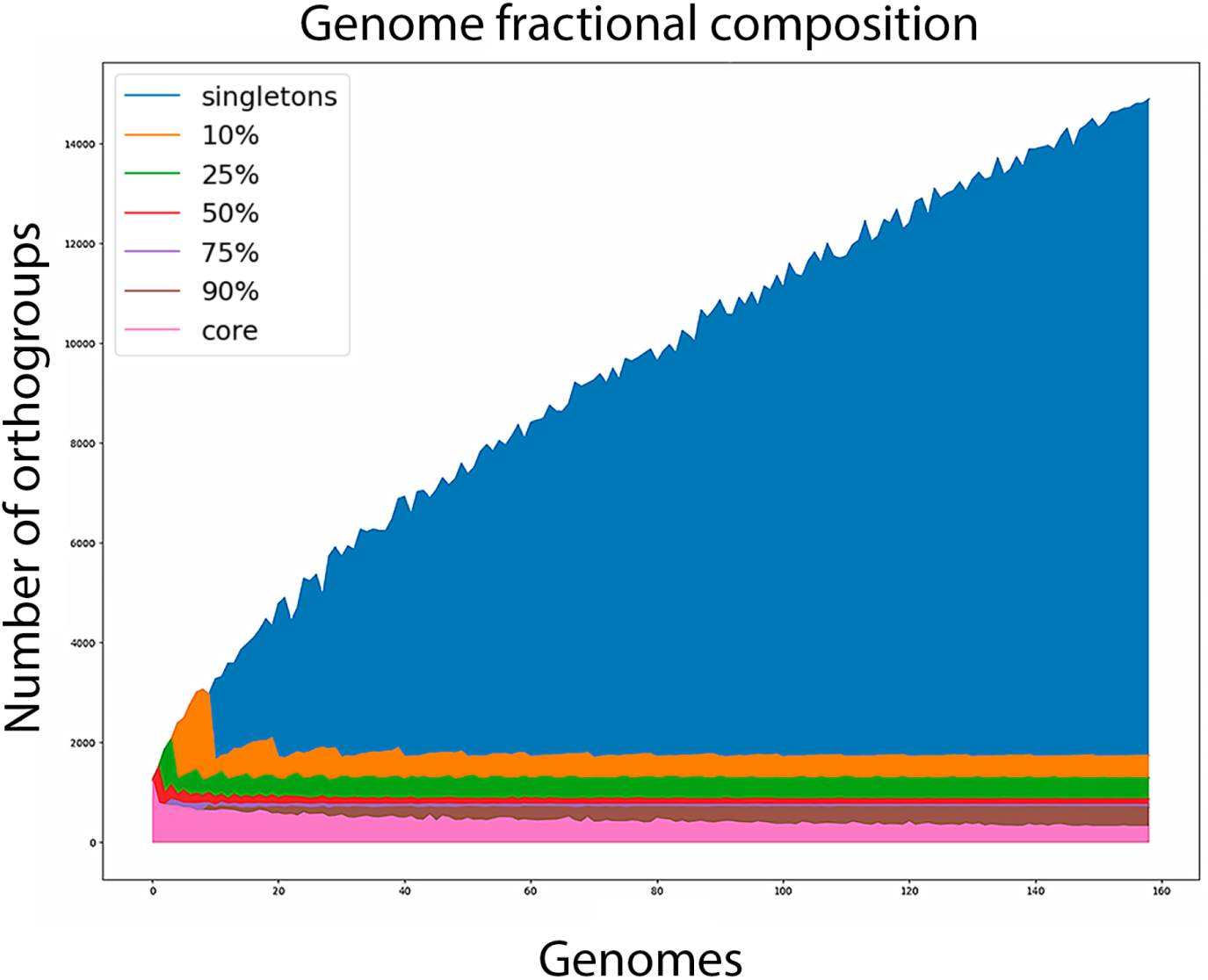
Number of genes present in a given fraction of *Wolbachia* genomes as dependent on the number of considered genomes. The topmost boundary shows the pan-genome size, the lowest boundary shows the core genome size, and the remaining boundaries show the percentile pan-genome sizes.

**Figure S3:**
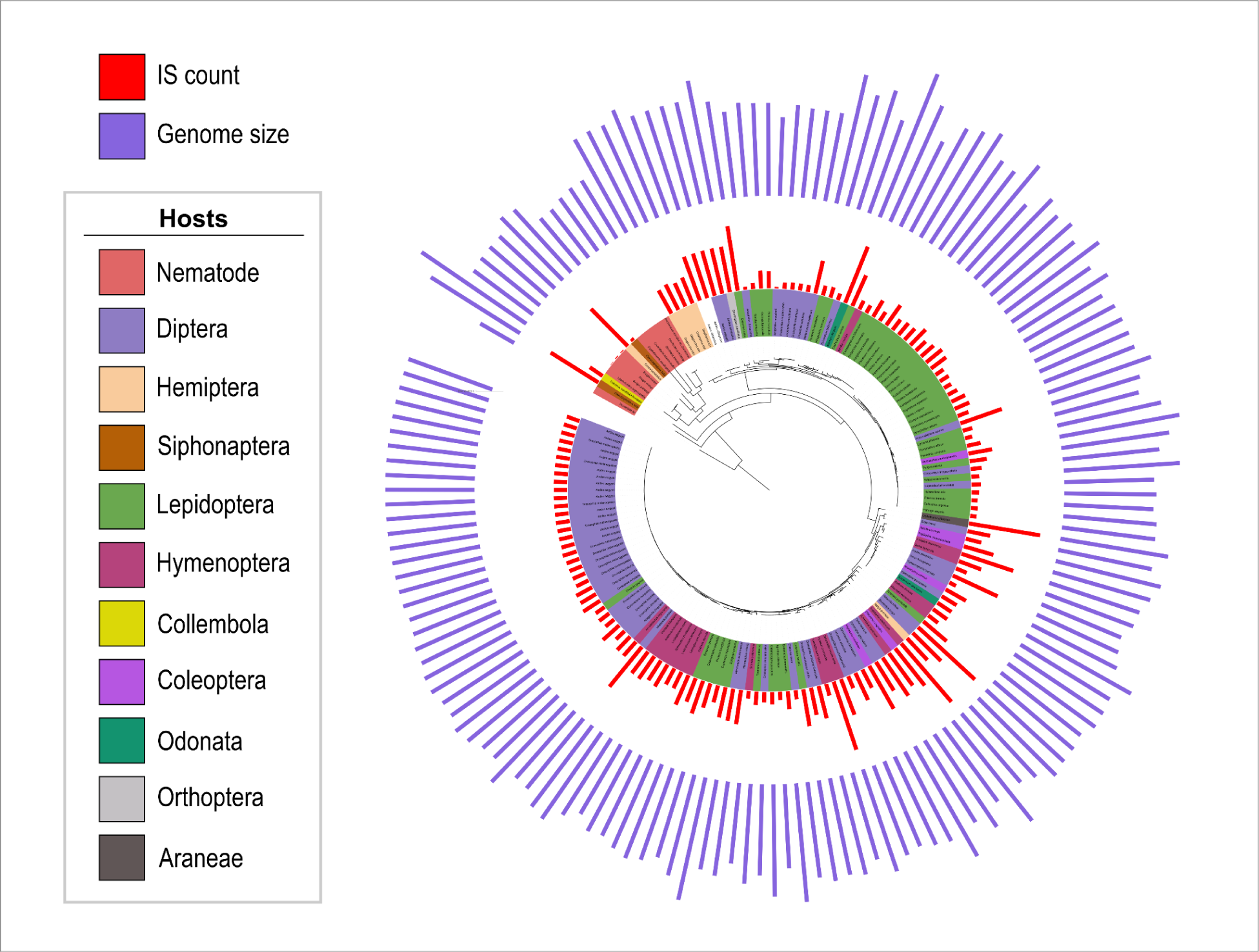
The number of ISs and the genome size shown on a phylogenetic tree.

**Figure S4:**
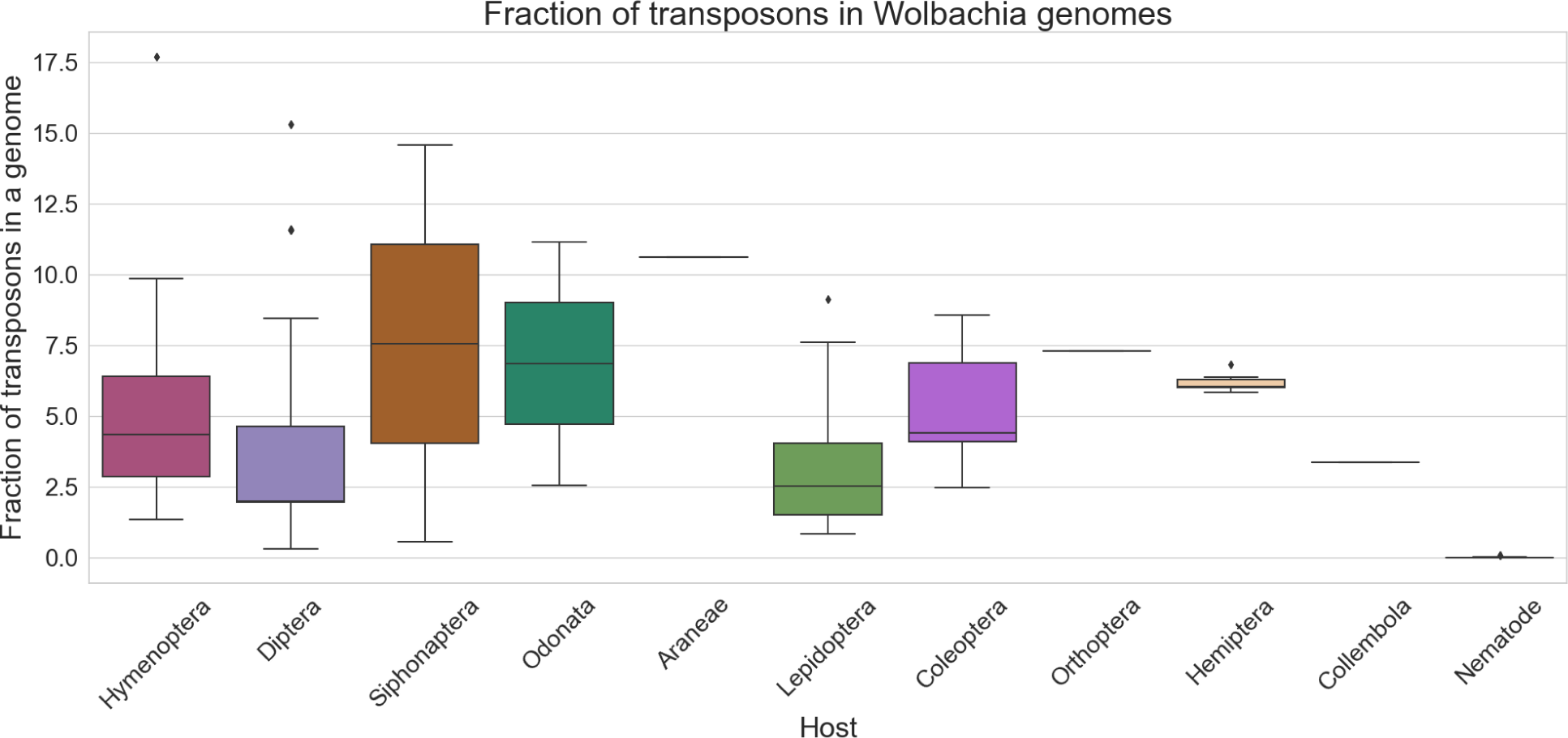
Fraction of transposons in *Wolbachia* genomes from different hosts.

**Figure S5:**
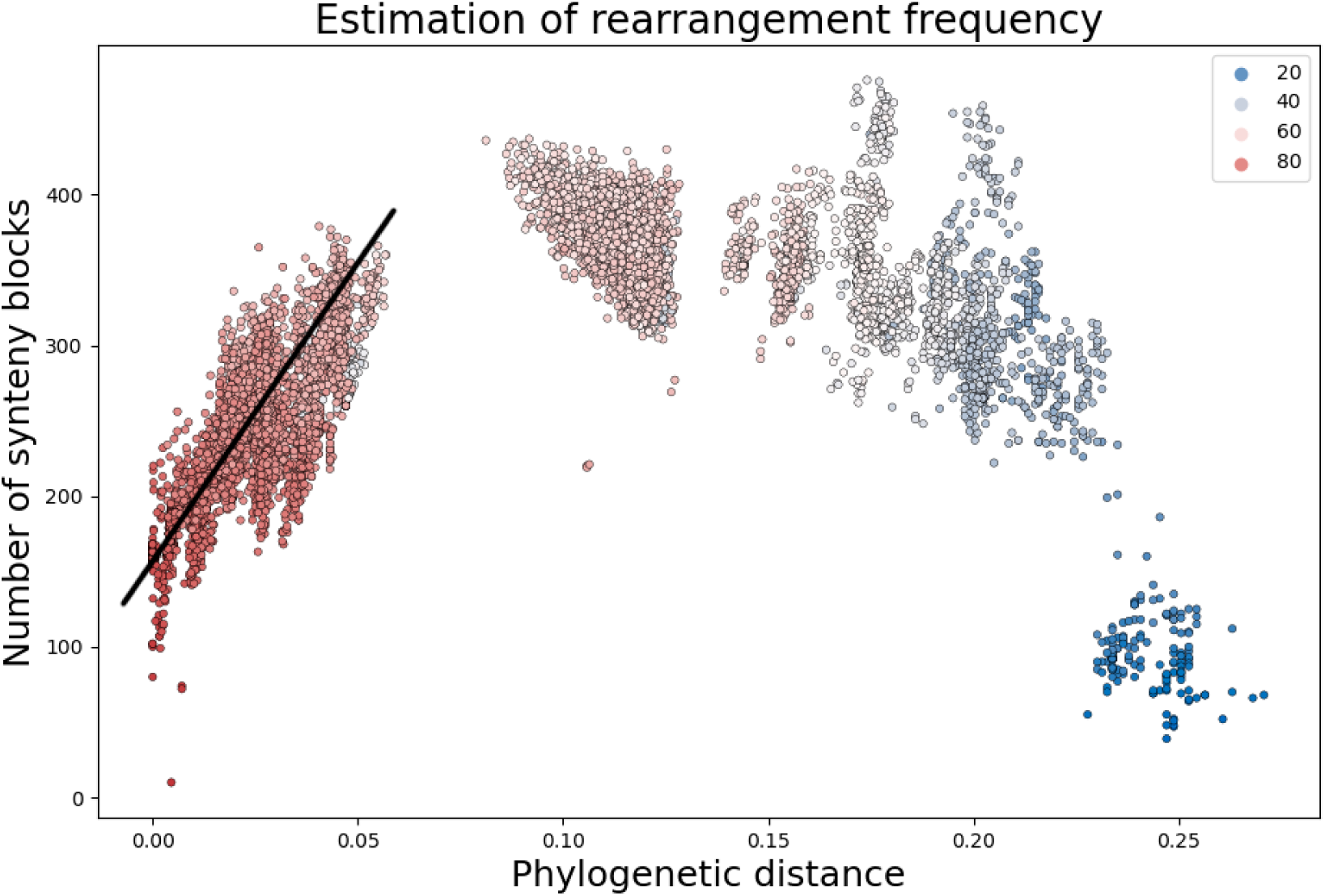
The number of locally collinear blocks and the distance between the genomes. Linear approximation of the relationships between these values at evolutionary distances <0.06 substitutions per site is shown by the black line.

**Figure S6:**
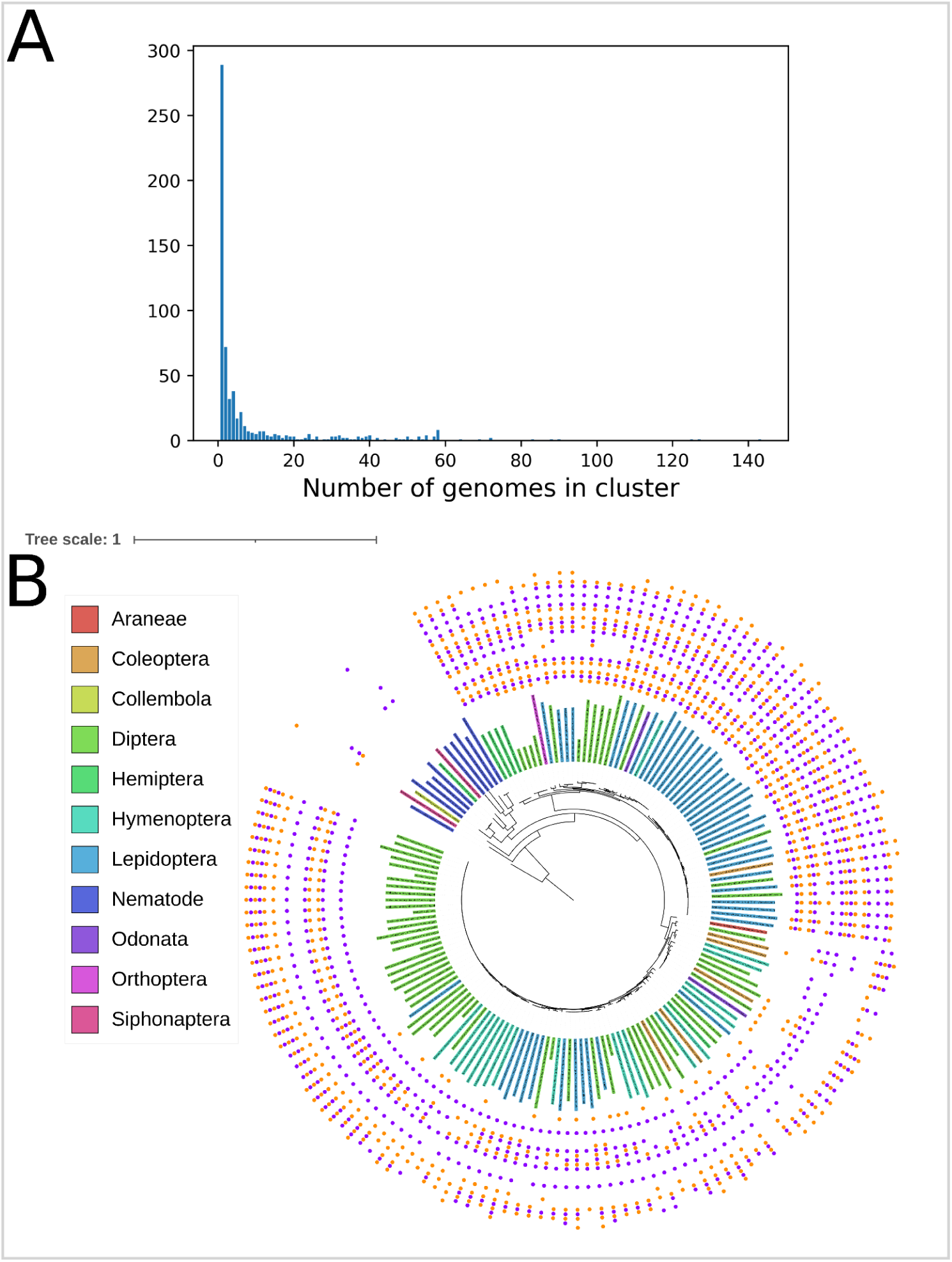
Occurrence of orthogroups containing ANK genes.

**Figure S7:**
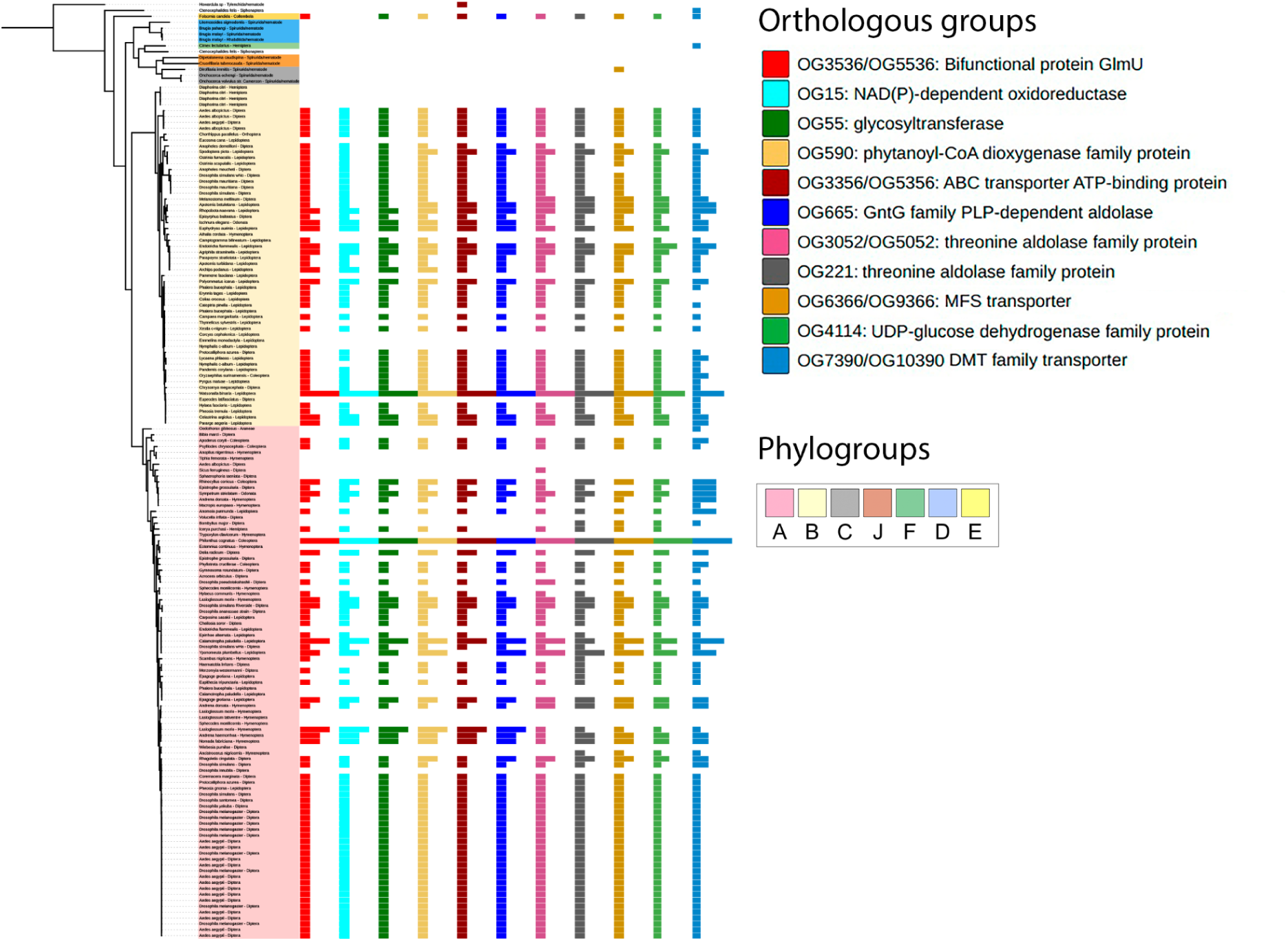
Segmental duplication in *Wolbachia* with the highest copy number variation. (A) Copy number of genes forming the cassette, shown in the *Wolbachia* phylogenetic tree. (B) Gene tree of UDP-glucose 6-dehydrogenase family protein gene, the member of gene cassette, revealing the cases of tandem duplication and GHT.

## References

Balvín O, Roth S, Talbot B, Reinhardt K. 2018. Co-speciation in bedbug Wolbachia parallel the pattern in nematode hosts. Scientific reports 8.

Bochkareva OO, Moroz EV, Davydov II, Gelfand MS. 2018. Genome rearrangements and selection in multi-chromosome bacteria Burkholderia spp. BMC genomics 19: 1–17.

Bordenstein SR, Bordenstein SR. 2022. Widespread phages of endosymbionts: Phage WO genomics and the proposed taxonomic classification of Symbioviridae. PLoS genetics 18: e1010227.

Brandis G, Hughes D. 2020. The SNAP hypothesis: Chromosomal rearrangements could emerge from positive Selection during Niche Adaptation. PLoS genetics 16.

Cabanettes F, Klopp C. 2018. D-GENIES: dot plot large genomes in an interactive, efficient and simple way. PeerJ 6: e4958.

Chafee ME, Funk DJ, Harrison RG, Bordenstein SR. 2010. Lateral Phage Transfer in Obligate Intracellular Bacteria (Wolbachia): Verification from Natural Populations. Molecular biology and evolution 27: 501.

Cooper BS, Vanderpool D, Conner WR, Matute DR, Turelli M. 2019. Wolbachia Acquisition by Drosophila yakuba-Clade Hosts and Transfer of Incompatibility Loci Between Distantly Related Wolbachia. Genetics 212.

Cordaux R, Pichon S, Ling A, Pérez P, Delaunay C, Vavre F, Bouchon D, Grève P. 2008. Intense transpositional activity of insertion sequences in an ancient obligate endosymbiont. Molecular biology and evolution 25.

Cui L, Neoh HM, Iwamoto A, Hiramatsu K. 2012. Coordinated phenotype switching with large-scale chromosome flip-flop inversion observed in bacteria. Proceedings of the National Academy of Sciences of the United States of America 109.

Darby AC, Armstrong SD, Bah GS, Kaur G, Hughes MA, Kay SM, Koldkjær P, Rainbow L, Radford AD, Blaxter ML, Tanya VN, Trees AJ, Cordaux R, Wastling JM, Makepeace BL. 2012. Analysis of gene expression from the Wolbachia genome of a filarial nematode supports both metabolic and defensive roles within the symbiosis. Genome research 22.

Darling AE, Miklós I, Ragan MA. 2008. Dynamics of genome rearrangement in bacterial populations. PLoS genetics 4.

Dedeine F, Vavre F, Fleury F, Loppin B, Hochberg ME, Bouletreau M. 2001. Removing symbiotic Wolbachia bacteria specifically inhibits oogenesis in a parasitic wasp. Proceedings of the National Academy of Sciences of the United States of America 98.

Dobrindt U, Chowdary MG, Krumbholz G, Hacker J. 2010. Genome dynamics and its impact on evolution of Escherichia coli. Medical microbiology and immunology 199.

Driscoll TP, Verhoeve VI, Brockway C, Shrewsberry DL, Plumer M, Sevdalis SE, Beckmann JF, Krueger LM, Macaluso KR, Azad AF, Gillespie JJ. 2020. Evolution of Wolbachia mutualism and reproductive parasitism: insight from two novel strains that co-infect cat fleas. PeerJ 8.

Edgar RC. 2004. MUSCLE: multiple sequence alignment with high accuracy and high throughput. Nucleic acids research 32: 1792.

Eugénio AT, Marialva MSP, Beldade P. 2023. Effects of Wolbachia on Transposable Element Expression Vary Between Drosophila melanogaster Host Genotypes. Genome biology and evolution 15: evad036.

Finn RD, Bateman A, Clements J, Coggill P, Eberhardt RY, Eddy SR, Heger A, Hetherington K, Holm L, Mistry J, Sonnhammer EL, Tate J, Punta M. 2014. Pfam: the protein families database. Nucleic acids research 42.

Foster J, Ganatra M, Kamal I, Ware J, Makarova K, Ivanova N, Bhattacharyya A, Kapatral V, Kumar S, Posfai J, Vincze T, Ingram J, Moran L, Lapidus A, Omelchenko M, Kyrpides N, Ghedin E, Wang S, Goltsman E, Joukov V, Ostrovskaya O, Tsukerman K, Mazur M, Comb D, Koonin E, Slatko B. 2005. The Wolbachia genome of Brugia malayi: endosymbiont evolution within a human pathogenic nematode. PLoS biology 3.

Hosokawa T, Koga R, Kikuchi Y, Meng XY, Fukatsu T. 2010. Wolbachia as a bacteriocyte-associated nutritional mutualist. Proceedings of the National Academy of Sciences of the United States of America 107: 769–774.

Hyatt D, Chen G-L, LoCascio PF, Land ML, Larimer FW, Hauser LJ. 2010. Prodigal: prokaryotic gene recognition and translation initiation site identification. BMC bioinformatics 11: 119.

Iturbe-Ormaetxe I, Burke GR, Riegler M, O’Neill SL. 2005. Distribution, expression, and motif variability of ankyrin domain genes in Wolbachia pipientis. Journal of bacteriology 187.

Katoh K, Standley DM. 2013. MAFFT Multiple Sequence Alignment Software Version 7: Improvements in Performance and Usability. Molecular biology and evolution 30: 772.

Kaur R, Siozios S, Miller WJ, Rota-Stabelli O. 2017. Insertion sequence polymorphism and genomic rearrangements uncover hidden Wolbachia diversity in Drosophila suzukii and D. subpulchrella. Scientific reports 7: 1–11.

Kaur R, Shropshire JD, Cross KL, Leigh B, Mansueto AJ, Stewart V, Bordenstein SR, Bordenstein SR. 2021. Living in the endosymbiotic world of Wolbachia: A centennial review. Cell host & microbe 29: 879–893.

Kojic M, Jovcic B, Miljkovic M, Novovic K, Begovic J, Studholme DJ. 2020. Large-scale chromosome flip-flop reversible inversion mediates phenotypic switching of expression of antibiotic resistance in lactococci. Microbiological research 241.

Laidoudi Y, Marie JL, Tahir D, Watier-Grillot S, Mediannikov O, Davoust B. 2020. Detection of Canine Vector-Borne Filariasis and Their Wolbachia Endosymbionts in French Guiana. Microorganisms 8.

Lechner M, Findeiss S, Steiner L, Marz M, Stadler PF, Prohaska SJ. 2011. Proteinortho: detection of (co-)orthologs in large-scale analysis. BMC bioinformatics 12.

Leclercq S, Giraud I, Cordaux R. 2011. Remarkable abundance and evolution of mobile group II introns in Wolbachia bacterial endosymbionts. Molecular biology and evolution 28.

Lefoulon E, Clark T, Borveto F, Perriat-Sanguinet M, Moulia C, Slatko BE, Gavotte L. 2020. Pseudoscorpion Wolbachia symbionts: diversity and evidence for a new supergroup S. BMC microbiology 20.

Letunic I, Bork P. 2021. Interactive Tree Of Life (iTOL) v5: an online tool for phylogenetic tree display and annotation. Nucleic acids research 49.

Liu B, Ren YS, Su CY, Abe Y, Zhu DH. 2023. Pangenomic analysis of Wolbachia provides insight into the evolution of host adaptation and cytoplasmic incompatibility factor genes. Frontiers in microbiology 14.

Lo N, Casiraghi M, Salati E, Bazzocchi C, Bandi C. 2002. How Many Wolbachia Supergroups Exist? Molecular biology and evolution 19: 341–346.

Mamirova L, Popadin K, Gelfand MS. 2007. Purifying selection in mitochondria, free-living and obligate intracellular proteobacteria. BMC evolutionary biology 7.

Manoj RRS, Latrofa MS, Epis S, Otranto D. 2021. Wolbachia: endosymbiont of onchocercid nematodes and their vectors. Parasites & vectors 14: 1–24.

Mendes FK, Vanderpool D, Fulton B, Hahn MW. 2021. CAFE 5 models variation in evolutionary rates among gene families. Bioinformatics 36.

Minh BQ, Schmidt HA, Chernomor O, Schrempf D, Woodhams MD, von Haeseler A, Lanfear R. 2020. IQ-TREE 2: New Models and Efficient Methods for Phylogenetic Inference in the Genomic Era. Molecular biology and evolution 37: 1530–1534.

Minkin I, Medvedev P. 2020. Scalable multiple whole-genome alignment and locally collinear block construction with SibeliaZ. Nature communications 11: 1–11.

Mosavi LK, Cammett TJ, Desrosiers DC, Peng Z-Y. 2004. The ankyrin repeat as molecular architecture for protein recognition. Protein science: a publication of the Protein Society 13: 1435.

Newton ILG, Rice DW. 2020. The Jekyll and Hyde Symbiont: Could Wolbachia Be a Nutritional Mutualist? Journal of bacteriology 202.

Nikoh N, Hosokawa T, Moriyama M, Oshima K, Hattori M, Fukatsu T. 2014. Evolutionary origin of insect-Wolbachia nutritional mutualism. Proceedings of the National Academy of Sciences of the United States of America 111.

Perrin A, Rocha EPC. 2021. PanACoTA: a modular tool for massive microbial comparative genomics. NAR Genomics and Bioinformatics 3: lqaa106.

Scholz M, Albanese D, Tuohy K, Donati C, Segata N, Rota-Stabelli O. 2020. Large scale genome reconstructions illuminate Wolbachia evolution. Nature communications 11: 1–11.

Seemann T. 2014. Prokka: rapid prokaryotic genome annotation. Bioinformatics 30.

Seferbekova Z, Zabelkin A, Yakovleva Y, Afasizhev R, Dranenko NO, Alexeev N, Gelfand MS, Bochkareva OO. 2021. High Rates of Genome Rearrangements and Pathogenicity of Shigella spp. Frontiers in microbiology 12: 628622.

Shelyakin PV, Bochkareva OO, Karan AA, Gelfand MS. 2019. Micro-evolution of three Streptococcus species: selection, antigenic variation, and horizontal gene inflow. BMC evolutionary biology 19: 1–15.

Siguier P, Perochon J, Lestrade L, Mahillon J, Chandler M. 2006. ISfinder: the reference centre for bacterial insertion sequences. Nucleic acids research 34.

Siozios S, Ioannidis P, Klasson L, Andersson SGE, Braig HR, Bourtzis K. 2013. The Diversity and Evolution of Wolbachia Ankyrin Repeat Domain Genes. PloS one 8.

Taylor MJ, Bordenstein SR, Slatko B. 2018. Microbe Profile: Wolbachia: a sex selector, a viral protector and a target to treat filarial nematodes. Microbiology 164.

Weinert LA, Araujo-Jnr EV, Ahmed MZ, Welch JJ. 2015. The incidence of bacterial endosymbionts in terrestrial arthropods. Proceedings. Biological sciences / The Royal Society 282.

Werren JH. 1997. Biology of Wolbachia. Annual review of entomology 42.

Xie Z, Tang H. 2017. ISEScan: automated identification of insertion sequence elements in prokaryotic genomes. Bioinformatics 33.

Zabelkin A, Yakovleva Y, Bochkareva O, Alexeev N. 2022. PaReBrick: PArallel REarrangements and BReaks identification toolkit. Bioinformatics 38.

Zabelkin A & Bochkareva O. BADLON: linear collinear Blocks Analysis: Distribution, Location, and compositiON. In preparation

Zug R, Hammerstein P. 2012. Still a Host of Hosts for Wolbachia: Analysis of Recent Data Suggests That 40% of Terrestrial Arthropod Species Are Infected. PloS one 7: e38544.

